# Risk factors, patterns and seropositivity of inter-epizootic Rift Valley fever virus of cattle in northern Tanzania

**DOI:** 10.1101/2025.10.02.679424

**Authors:** James A. Mlacha, Athumani M. Lupindu, Robinson H. Mdegela, Oliva K. Manangwa, Furaha Mramba, Jessica Clark, Paul Johnson, Sarah Cleaveland, Jennifer S. Lords

## Abstract

1.1

**Background:** Rift Valley Fever (RVF) is a zoonotic viral disease affecting both animals and humans, with significant public health and economic implications. In Tanzania, a series of RVF outbreaks in livestock have been reported. The epidemiological information about hosts’ exposures status, time, location and risk factors that contribute to virus maintenance between one epizootic and the other, is important for preparedness and intervention plans. This study investigated the seroprevalence and potential determinants of RVFV exposure in cattle in Babati, Hai, and Moshi Districts between the last RVF outbreak in 2007 and the anticipated El Niño rains of 2024, which could precipitate the next outbreak.

**Methods:** A total of 1,627 cattle serum samples (790 from Babati, 518 from Moshi, and 319 from Hai), were collected between August and October 2023 from both urban and rural areas, were analyzed using a competitive ELISA to detect RVFV-specific antibodies. Data on cattle biodata, management practices, environmental factors, were obtained through structured questionnaires and field records. Sero-prevalence was determined by descriptive statistics, while risk factors were quantified by multivariable logistic regression and RVF seropositivity was plotted on administrative boundaries of Babati, Moshi, and Hai districts by using QGIS.

**Results:** The overall RVFV seroprevalence was 15.0% (95% CI: 13.26 – 16.73). Seropositivity was 22.2% (95% CI: 18.62-25.78) in Moshi, 12.2% (95% CI: 8.63-15.82) in Hai and 11.4% (95% CI 9.18-13.61) in Babati. Age (OR = 1.22, 95% CI: 1.15–1.3, p<0.001), sex (OR = 0.53, 95% CI: 0.36–0.79, *p* = 0.0017), herd size (OR=0.2201; 95% CI: 0.095–0.5101; p=0.0004), water access along irrigated crop fields (OR=3.7181; 95% CI: 1.5868–8.7121; p=0.0025), and geographical location (OR = 2.23, 95% CI: 1.63–3.06, *p* < 0.001) were significant predictors of seropositivity. Spatial visualization revealed clustering of seropositive herds in villages near lake Manyara and Tarangire National parks in Manyara region, also seropositive clusters in villages near irrigated areas, Nyumba ya Mungu dam and floodplains in lower Moshi.

**Conclusion:** This study highlights ongoing inter-epizootic exposure of cattle to RVFV in northern Tanzania. The RVFV is not randomly distributed during inter-epizootic periods, but rather forms seropositive clusters. Most significant determinants of seropositivity includes age, sex, herd size, water access along irrigated crop fields and geographical location Moshi showing higher seropositivity.

## 1.2 INTRODUCTION

Rift Valley fever (RVF) is a mosquito-borne zoonotic disease affecting both humans and livestock, caused by Rift Valley fever virus (RVFV), an enveloped, negative-sense single-stranded RNA virus of the genus *Phlebovirus* (Pepin *et al*., 2010). In livestock RVF is characterized by abortion, fever, nasal discharges, ocular discharges, hemorrhagic diarrhea and higher mortality rate especially in young animals (Wensman *et al*., 2015). In humans, the disease ranges from mild febrile illness to severe hemorrhagic manifestations (Rwegoshola *et al*., 2023), malnutrition and high case fatality rates in severe cases (Ahmed *et al*., 2018) Patterns of Rift Valley fever virus (RVFV) occurrence are strongly associated with seasonal rainfall, flooding, and vegetation dynamics, which create ideal breeding habitats for mosquito vectors (Anyamba *et al*., 2010). Multiple mosquito species involved in RVFV transmission, are *Aedes* spp. serving as primary reservoirs capable of maintaining the virus vertically in eggs during dry periods, while *Mansonia* spp, *Culex* spp. and *Anopheles* spp. act as secondary amplifiers during epizootics (Lumley *et al*., 2017).

Tanzania shows a clear temporal pattern, with major outbreaks occurring after every 5 to 20 years (Mohamed et al., 2010), while spatially, transmission is concentrated in northern and central regions where high livestock density and suitable mosquito habitats coincide (Sindato et al., 2014; Sang et al., 2016). The first outbreak was recorded on 1930, then several periodic outbreaks occurred in 1947, 1957, 1977 and 1997 (Wensman *et al*., 2015). The last outbreak of RVF occurred in 2006/2007 and was reported in Manyara, Tanga, Dodoma, Morogoro, Dar es Salaam, Coast, Iringa, Mwanza and Singida regions of Tanzania (Kifaro *et al*., 2014). It resulted to financial loss due to declining of livestock export by 54%, resulting in an estimated loss of $352,750. In addition, livestock mortality cattle deaths valued at approximately $4,243,250, and losses from goats and sheep at $2,202,467(Sindato e*t al*., 2012), this loss negatively impacted national economies (Ahmed *et al*., 2018).

Determinants of infection vary across hosts and ecosystems, with factors such as age, sex, geographic location, herd density, and proximity to irrigation or flood-prone areas influencing seroprevalence in livestock and humans (LaBeaud *et al*., 2011). RVF is typically assumed to have two events which are epizootic event and inter-epizootic also known as enzootic event. Epizootic cycle is define as years in which RVF outbreaks were reported in at least three districts (Murithi *et al*., 2011). Findings from (Linthicum *et al*., 1985) demonstrated that RVFV can be vertically transmitted in Aedes mosquitoes, with infected females passing the virus to their offspring, and led to the assumption that, between major epizootics, RVFV transmission exclusively occurs through this vertical route. Multiple studies have, however, demonstrated that RVFV actively circulates during inter-epizootic periods, with serological and molecular evidence of infection detected in livestock and humans outside recognized outbreaks (LaBeaud *et al*., 2008; Sumaye e*t al*., 2013). Despite inter-epizootic transmission with reservoir maintenance, as earlier reported (De Glanville *et al*., 2022), little is known about what is driving the maintenance of transmission to livestock during inter-epizootic period.

Over the last two decades East Africa has undergone substantial transformations that could alter RVFV transmission dynamics, including rising livestock densities, increased peri-urban livestock keeping, and expansion of crop irrigation. These changes have raised host availability to the RVFV, increase human–animal contact, and potentially created new mosquito habitats, all of which can favor RVFV persistence and outbreak initiation (Sindato *et al*., 2015).

Differences in livestock management and environmental conditions between rural and peri-urban areas may influence RVF exposure. Rural areas typically have larger herds, more diverse livestock species, and greater access to natural water sources, whereas peri-urban areas often have smaller herds, limited species diversity, and water sources concentrated around human settlements. These variations could affect mosquito breeding habitats and contact rates between vectors and hosts. It is posited that RVFV determinants and seroprevalence in cattle varies between rural and peri-urban areas during inter-epizootic. This study explains RVF seroprevalence, spatial distribution and determinants of RVF during inter-epizootic period which will help in development of proper policy on control and prevention strategies hence breaking the epizootic cycles.

## 1.3 MATERIALS AND METHODS

### 1.3.1 Study Area

This study was conducted in one district from Manyara Region (Babati district) due to its large area coverage and two districts from Kilimanjaro Region (Hai and Moshi Rural Districts). These areas were selected due to their ecological and economic importance in livestock farming, previous RVF outbreaks, and environmental conditions that favor mosquito breeding, high density of livestock (cattle, sheep, goats) and wildlife populations (**Figure 1**).

**Figure 1.**
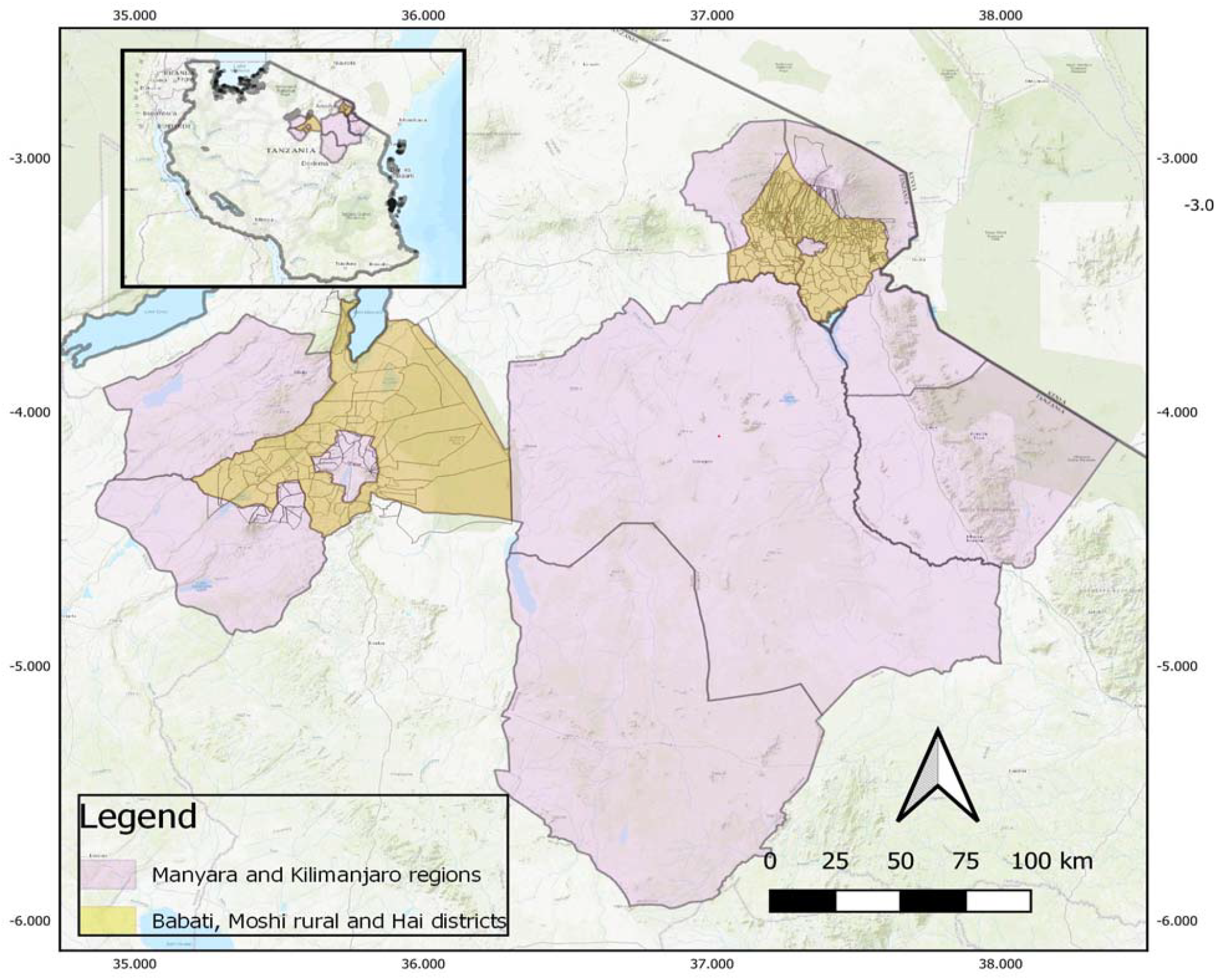
A map showing Babati, Moshi rural and Hai Districts which are in Northern part of Tanzania (created using QGIS).

Kilimanjaro and Manyara regions are situated in Northern part of Tanzania, and especially lower Moshi there are numerous water streams for irrigation on rice and sugar cane farms which cover an area of about 1100 hectares (Rwegoshola *et al*., 2023). The area has two rainy seasons which are the long rains seasons from March to May and the short rainy season from October to December. The average rainfall is 900 mm (Rwegoshola *et al*., 2023).

Babati district, in Manyara region is located in the eastern wing of the Rift Valley, which was most affected in the previous outbreaks including the last one during 2006/2007 (Salekwa et al., 2019). Geographically, Babati District lies approximately between latitudes 3°00’ and 4°30’ South and longitudes 35°00’ and 36°00’ East. Babati experiences average annual temperatures ranging from 16°C to 30°C and rainfall averages between 500 mm and 1,200 mm per year (Babati district council, 2022).

Hai and Moshi Rural Districts from Kilimanjaro, are situated between latitudes 3°00’ and 3°30’ South and longitudes 37°00’ and 37°45’ East. The climatic conditions in these areas vary slightly due to differences in altitude and proximity to Mount Kilimanjaro. Temperature range between 15°C and 28°C., and there is slightly higher annual rainfall, relative humidity typically ranging from 55% to 75%, depending on the season (Moshi district council; Hai district council, 2022).

According to the 2022 Tanzania Population and Housing Census, Babati District has a population of approximately 420,479, comprising 206,487 males (49.1%) and 213,992 females (50.9%). Hai District has a population of 274,498 with 131,951 males (48.1%) and 142,547 females (51.9%), while Moshi Rural District accounts for 466,740 people, including 224,218 males (48.0%) and 242,522 females (52.0%). Livestock production forms a significant part of rural livelihoods in all study areas. Babati District which covers 5,608 km^2^ is home to approximately 259,717 cattle, 179,653 goats, 76,086 sheep, 7,765 donkey, 230,440 chicken, 7,968 pigs and 17,278 dogs. In Kilimanjaro Region, Hai District covers 1,011 km^2^ holds about 100,751 cattle, 34,357 sheep; 16,820 pigs, 58,838 goats and 225,513 chickens while Moshi Rural District with 1,713 km^2^ coverage keeps an estimated 141,796 cattle, 105,780 goats, and 34,187 sheep, 53,879 pigs, 24,167 dogs and 1,309,985 chickens (Moshi District Council, 2022; Hai District Council, 2022; Babati District Council, 2023) The Agro-pastoral and mixed farming systems practiced in these regions underscore the importance of livestock in both economic and social contexts.

### 1.3.2 Study Design and sample size estimation

We carried out a cross-sectional study of cattle in rural and peri-urban areas of Manyara (Babati district) and Kilimanjaro region (Hai and Moshi rural) between August to October 2023. Blood sera were collected from 38 villages across all three districts.

Sample size determination was carried out using the R package Generalized linear mixed models GLMMmisc (Johnson *et al*., 2015), which allows simulation-based power calculations for complex hierarchical designs. Simulated data were generated to reflect livestock seropositivity structured by household and village, with wards stratified as high- or low-risk areas. Logistic regression models were fitted to the simulated datasets to evaluate the ability to detect differences in seroprevalence between hypothesized high- and low-risk wards. Power was defined as the proportion of simulations that successfully identified a statistically significant difference in risk taking into considerations the cattle populations in all three districts.

Different sample sizes were tested in the simulations to identify the number of animals required for robust detection of the expected effect size. Based on these simulations, sampling 1,400 cattle (ten animals from four households across 35 villages) provided 87% power to detect a three-fold difference in RVFV risk between high- and low-risk wards. Villages were identified in Kilimanjaro and Manyara regions and later they were selected from Babati, Hai and Moshi rural districts. From each village, households keeping cattle were randomly selected and categorized as rural or peri-urban using estimates from google building density and population density.

### 1.3.3 Farm and individual animal selection

Blood sample collection followed a multi-stage random selection process to ensure representativeness and to minimize bias. From each village, households keeping cattle were randomly chosen. Finally, from each household, up to 10 animals were sampled in extensive systems. In peri-urban areas where households kept fewer than 10 cattle, all animals were included

### 1.3.4 Data Collection

### 1.3.4.1 Source of data

Total of 1627 blood samples (790 from Babati, 518 from Moshi and 319 from Hai), aseptically collected between August and October 2023 from cattle above six months to obtain blood serum. Age, sex, breed, vaccination history, farming practice, size of the herd and settlement type (rural or peri-urban) was collected for the purpose of assessing determinants of RVFV seropositivity. Geographical locations were recorded using GPS function embedded in Open Data Kit (ODK) Toolbox on Android tablet for spatial mapping of the disease distribution in respect of RVF virus infection (Hartung et al., 2010)

#### 1.3.4.2 Administration of questionnaire

To assess potential risk factors for herd-level RVFV seropositivity, a structured questionnaire was administered to livestock keepers at the time of sample collection. From the total 206 sampled households, 153 were randomly selected to participate in the survey. The initial goal was to include four households per village. However, in peri-urban villages where herd sizes were small, more than four households had to be sampled in order to reach the minimum target of 40 cattle per village. In such cases, households were still selected randomly for inclusion in the questionnaire administration. Primary data on herd composition, livestock management, herd size, use of acaricides and insecticides, vaccination history, water access, grazing practices, year-round animal presence, and introduction of new animals were collected through face-to-face interviews using a structured questionnaire

#### 1.3.4.3 Blood sample collection and serum preparation

Blood samples were aseptically drawn from the jugular vein using sterile vacutainer needles and plain vacutainer tubes (without anticoagulant). Each animal was properly restrained, and the venipuncture site was disinfected, 8-10ml blood samples were drawn and put in a cool box with ice packs before separating the serum from whole coagulated blood by centrifugation at 1500g for ten minutes. The sera were transferred into labeled 1.8 ml cryovials and kept at –20°C for short-term storage, then long term storage was done at −80 °C in Kilimanjaro Clinical Research Institute (KCRI). Later all samples collected from all three districts were then transported to Vector and Vector Borne Research Institute (VVBD/TVLA Tanga) and kept in a freezer at −80 °C until laboratory analysis.

During sample collection, animal that had no identifications were given ear tags with unique numbers. New identified animals and those which had their tags already from each household were sampled and important information like age, sex, breed, number of parities, herd size and management practices were obtained from animal owners, herdsmen or family members.

#### 1.3.4.4 Laboratory procedures

A total of 1,627 cattle serum samples were analyzed for antibodies against Rift Valley fever virus (RVFV) using a competitive enzyme-linked immunosorbent assay (cELISA) (ID Screen® Rift Valley Fever Competition Multi Species, IDvet, Grabels, France). Prior to testing, all serum samples were retrieved from -80°C storage and allowed to thaw at room temperature. Once fully thawed, the samples underwent heat inactivation at 56°C for 60 minutes to minimize any potential biohazard risk (Soltis *et al*., 1979). The ELISA procedure was then conducted strictly following the manufacturer’s instructions. Briefly, 50 µL of each sample was diluted with diluent to obtain 100 µL, and along with the provided positive and negative controls, were added in duplicate to the ELISA plate pre-coated with RVFV antigen. The plate was incubated at 37°C for 1 hour. After incubation, wells were washed three times using the wash buffer supplied in the kit to remove unbound antibodies. Subsequently, 100 µL of the conjugate solution was added to each well and the plate was incubated again at 21°C (room temperature) for 30 minutes. Another washing step followed, after which 100 µL of the substrate solution was added to each well. The plate was incubated for 15 minutes at room temperature in the dark, and then 100 µL of the stop solution was added to halt the reaction. Optical densities (OD) were read at 450 nm using an ELISA microplate reader as per manufacture’s instruction. Results were expressed as a percentage of the sample-to-negative control ratio (S/N%). Samples with an S/N% less or equal to 40% were considered positive for RVFV antibodies, as per the kit’s interpretive criteria (Bazanow et al., 2018). Throughout the procedure, quality control measures were strictly adhered to, and test validity was confirmed by the expected performance of the kit controls.

#### 1.3.4.5 Data analysis

Descriptive analysis was done using Microsoft Excel 2016 to handle the obtained data and provide an overview of the study population and Rift RVFV seropositivity patterns. The overall seroprevalence of RVFV in cattle was calculated as the proportion of animals testing positive by cELISA relative to the total number of animals tested. Seroprevalence estimates were accompanied by 95% confidence intervals to account for sampling variability.

To address whether the seroprevalence of RVFV exposure varies between rural and peri-urban settings, Chi-square (*χ*^2^) test was used to check for statistical differences between the two settings, as well as among the three study districts at a significance level of 0.05 (Hosmer et al., 2013)

Risk factors for RVFV seropositivity were done by two separate analyses on individual-animal level factors and herd-level management factors using multivariable logistic regression with backward stepwise elimination. The model was built after conducting univariate logistic analyses of four individual-animal level determinants and 13 herd-level determinants, using a liberal inclusion threshold of *p* > 0.25 as recommended by Hosmer et al. (2013). All analyses were performed in Epi Info™ 7.2.6.0 (2023). Interaction terms were included in the final model to check for statistical significance and the likelihood ratio test was used to test the goodness of fit of the model. At the individual-animal level, explanatory variables included host characteristics such as age, sex, settlement type (rural or peri-urban), and geographical location assessed in relation to the ELISA serostatus of each sampled animal.

At the herd-level analysis, a household was classified as seropositive if at least one animal tested positive by ELISA, and seronegative if none were positive. Herd-level explanatory variables included livestock management practices such as herd composition (species kept), herd size, use of acaricides and insecticides for mosquito control, vaccination history, access to water sources near crop fields, grazing on crops, year-round presence of animals, and introduction of new animals into the herd.

Mapping and spatial visualization was performed using QGIS version 3.32, with sampling locations plotted on administrative boundaries of Babati, Manyara, and Hai districts. Seroprevalence was visualized through color-coded point maps, which facilitated the identification of the spatial distribution of seropositive animals and highlighted isolated high-risk areas (QGIS Development Team, 2023

#### 1.3.4.6 Ethical consideration

This study was approved by Sokoine University of Agriculture with Reference No. SUA/DPRTC/MPV/D/2024/0015/03. Ethical Clearance Certificate was obtained from Tanzania Livestock Research Institute (TALIRI) with Reference No. TLRI/RCC.23/018 and from Commission for Science and Technology (COSTECH) with permit number 2022-750-NA-2022-149. Permission was sought to relevant local authorities in Kilimanjaro and Manyara regions while the informed written consent was obtained from all participants for voluntary participation.

## 1.4 RESULTS

### 1.4.1 Farms and study animals’ characteristics

Of the total of 1,627 cattle sera samples, 513 were male (31.5%) and 1,114 were female (68.5%), Age ranged from 6 months to 15 years, the animals aged two years or below were classified as young, comprising 35.89% (n = 584) of the total sample, while those older than two years were categorized as adults, accounting for 64.10% (n = 1,043) Cattle sera from Babati were 48.56% (n=790), Hai were 19.60% (n=319), and from Moshi Rural were 31.84% (n=518). Of the cattle sera collected, 62.26% (n = 1,013) were from rural settlements, while 37.74% (n = 614) were from peri-urban settlements.

### 1.4.2 Seroprevalence of Rift Valley Fever

The overall prevalence of RVF for Northern Tanzania (Kilimanjaro and Manyara) was 15.0% (95% CI: 13.26 – 16.73), and those from rural settlement was 15.60% (95% CI: 13.36 – 17.83), and from peri urban settlement was 14.01% (95% CI: 11.26 – 16.75) as shown in (**Table 1**). There was no statistically significant difference in RVFV seropositivity between cattle from rural and peri-urban settlement types (*χ*^*2*^ = 0.759, *p* = 0.424). Seroprevalence varied by district, the highest seroprevalence was observed in Moshi Rural, with 22.20% (95% CI: 18.62-25.78) of cattle testing positive, followed by Hai with 12.23% (95% CI: 8.63-15.82) and Babati with 11.39% (95% CI 9.18-13.61). A chi-square test revealed that these differences were statistically significant (*χ*^*2*^ *= 31*.*69, df = 2, p < 0*.*001*).

**Table 1:** Seroprevalence of Rift Valley Fever Based on individual-animal level variables.

| Variables | Category | Frequency | Positive | Prevalence | 95% CI |
| --- | --- | --- | --- | --- | --- |

|  |  |  |  | (%) |  |
| --- | --- | --- | --- | --- | --- |
| **District** | Moshi | 518 | 115 | 22.2 | 18.62-25.78 |
|  | Babati | 790 | 90 | 11.39 | 9.18-13.61 |
|  | Hai | 319 | 39 | 12.23 | 8.63-15.82 |
| **Sex** | Male | 513 | 36 | 7.02 | 4.81-9.23 |
|  | Female | 1114 | 208 | 18.67 | 16.38-20.96 |
| **Settlement types** | Rural | 1013 | 158 | 15.6 | 13.36-17.83 |
|  | Peri-urban | 614 | 86 | 14.1 | 11.26-16.75 |
| **Age** | Young | 584 | 42 | 7.19 | 5.1-9.29 |
|  | Adult | 1043 | 202 | 19.37 | 16.97-21.77 |

### 1.4.3 Spatial patterns of Rift Valley fever virus seropositivity

Spatial visualization indicated that, Seropositive cattle were not randomly distributed across the study area, but a clustering of RVFV exposure. In the Lower Moshi along the Nyumba ya Mungu corridor, villages adjacent to the Nyumba ya Mungu dam particularly Chemchem, Kochakindo and Mikocheni exhibited elevated levels of seropositive cattle compared to the other villages. Similarly, in irrigated agro-ecosystems, villages such as Rau River, Oria, Mawala and Kiterini, demonstrated large seropositive clusters (9-19 positive cattle) size, consistent with the creation of mosquito-friendly habitats around sugarcane and rice irrigation schemes **(Figure 2)**

**Figure 2.**
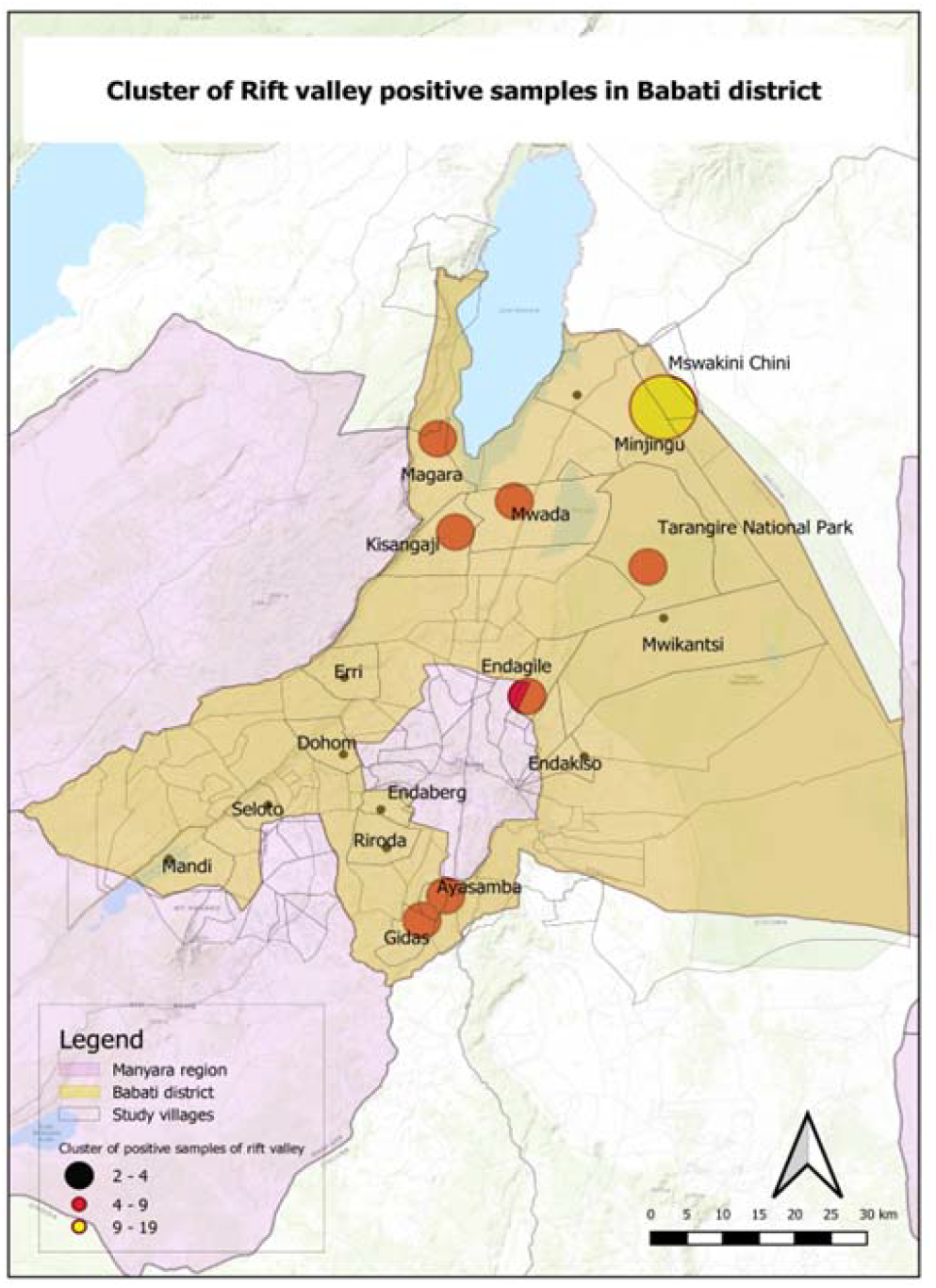
Spatial distribution of Rift Valley Fever positive animals in Babati Districts.

In Babati District, Minjingu and Kakoi villages located near Tarangire National Park showed large size of seropositive cluster compare to other villages. The wetlands areas specifically Mwada, Kisangaji and Manyara villages which are closer to Lake Manyara showed increased clustering of seropositive cattle compare to the rest of the villages, suggesting heightened risk at wildlife–livestock interfaces **(Figure 3)**.

**Figure 3.**
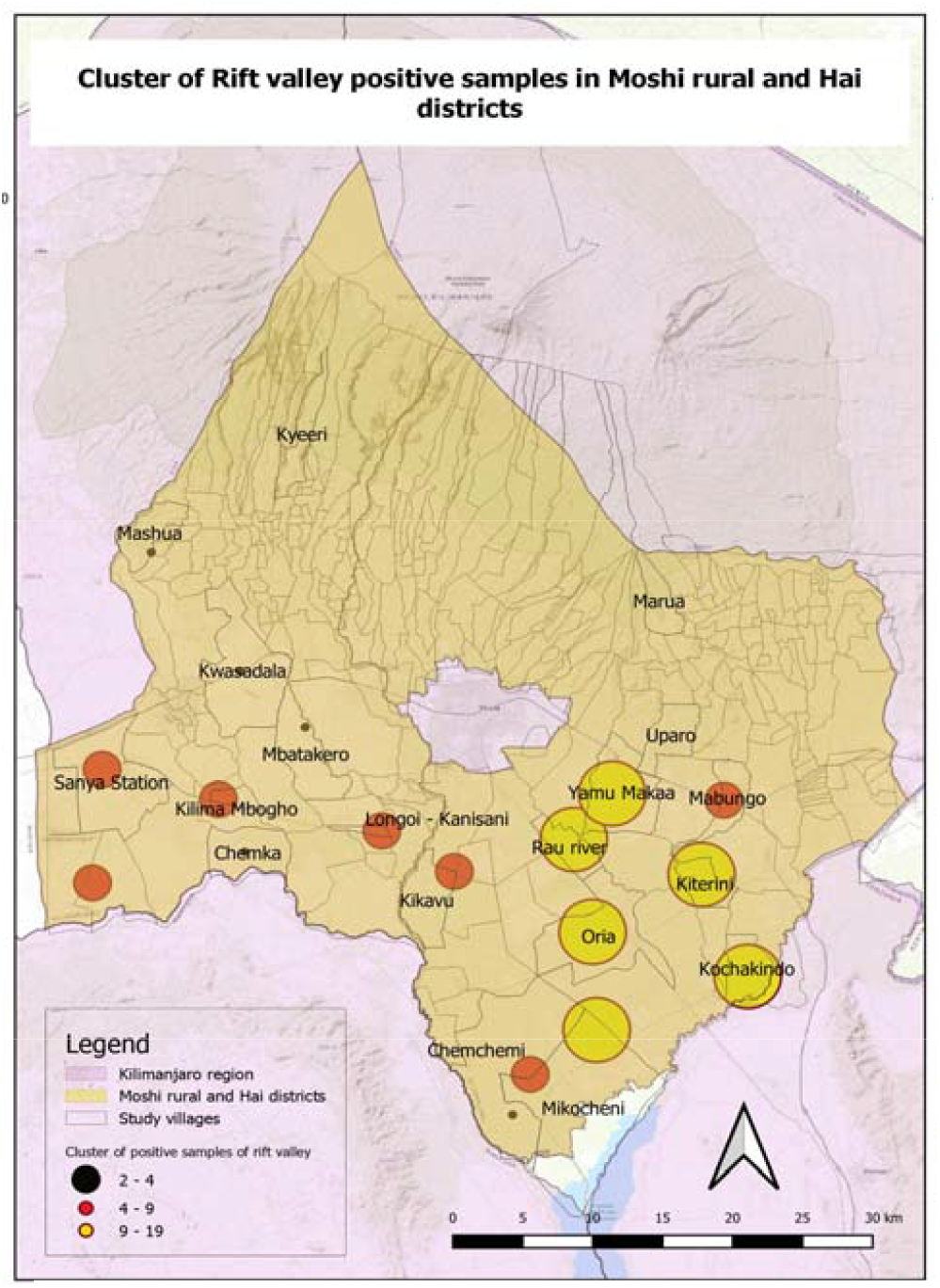
Spatial distribution of Rift Valley Fever positive animals in Hai and Moshi Districts.

### 1.4.4 Determinants for Rift Valley Fever virus exposure

Four individual-animal level variables (age, sex, settlement types and location) and seven herd-level variables (herd size category, year-round presence of cattle, grazing along the crops, accessing water along the irrigated crops, vaccination, use of insecticides to control mosquitoes and sharing shelters with other animals) were qualified for multivariable logistic regression.

The final model results of multivariable logistic regression analysis on individual-animal related determinants identified sex, age, and location as significant determinants of RVFV seropositivity in cattle. Male cattle had significantly lower odds of being seropositive compared to females (OR = 0.53, 95% CI: 0.36–0.79, *p* = 0.0017). Age was positively associated with seropositivity, with each unit increase in the age by one year the log odds of RVFV seropositivity increases by 0.1955 holding other variable constants, Coefficient (β= 0.1955, 95% CI 0.1385–0.2525, with *p* < 0.001). this analysis also concurred with that of chi-square test on cattle from Moshi Rural being two times more likely to be seropositive compared to those from Babati and Hai (OR = 2.23, 95% CI: 1.63–3.06, *p* < 0.001). These findings suggest that both age and sex as animal factors, and geographic location are important predictors of RVFV exposure.

While the final model for herd-level management related variables retained two predictors of RVFV Seropositivity. Herds categorized as small were approximately less likely to have RVFV seropositive cattle compared to large herds (OR=0.2201; 95% CI: 0.095–0.5101; p=0.0004), indicating that small herds had about 4 times lower odds of seropositivity. Livestock with access to water in crop fields had 3.7 times more odds of being seropositive than those without such access (OR=3.7181; 95% CI: 1.5868–8.7121; p=0.0025). This means that, large herd size and grazing animals around irrigated crops fields are herd-level related risk factors for RVFV Seropositivity as shown in **(Table 2)**:

**Table 2:** Risk factors associated with RVFV Seropositivity in cattle in Northern Tanzania.

| Variables | Category | Frequency | Positive | Prevalence (%) | OR | 95% CI |
| --- | --- | --- | --- | --- | --- | --- |
| Individual-animal level risk factors |  |  |  |  |  |  |
| District | Moshi | 518 | 115 | 22.2 | 2.2 | 1.63-3.06 |
|  | Babati | 790 | 90 | 11.39 |  |  |
|  | Hai | 319 | 39 | 12.23 |  |  |
| Sex | Male | 513 | 36 | 7.02 | 0.5 | 0.36-0.79 |
|  | Female | 1114 | 208 | 18.67 |  |  |
| Age | Adult | 1043 | 202 | 19.37 | 1.22 | 1.15-1.29 |
|  | Young | 584 | 42 | 7.19 |  |  |
| Herd-level Risk factors |  |  |  |  |  |  |
| Herd size category | Small | 98 | 48 | 48.98 | 0.22 | 0.01-0.51 |
|  | Large | 55 | 46 | 83.64 |  |  |
| Access water near crop field | Yes | 51 | 42 | 82.35 | 3.72 | 1.59-8.71 |
|  | No | 102 | 52 | 50.98 |  |  |

Both final model for host-related and management-related determinants demonstrated good fit, as indicated by the likelihood ratio test with (p < 0.001) confirming that the predictors collectively improved the model fit to the data.

## 1.5 DISCUSSION

The finding from this study provides further evidence of RVFV circulation during inter-epizootic period (IEP), indicating overall seroprevalence of 15.0% in Northern Tanzania. This prevalence suggests ongoing viral circulation of the virus, likely maintained through silent transmission cycles involving vectors and possibly wildlife reservoirs. Although no clinical outbreaks were reported during the sampling period, the detectable antibody levels confirm that RVFV continues to persist endemically within the ecosystem. It is higher from what was reported by previous study of 4.4% (De Glanville *et al*., 2022) but lower compared to 29.2% reported in Morogoro (Matiko et al., 2018).

This variation could be attributed by different sampling period, sample size, increased vector activities or increased livestock movement. High seroprevalence (22.2%) in Moshi Rural during an inter-epidemic period, with a highly significant difference among districts (χ^2^ = 31.06, *p* < 0.001), reinforces evidence of ongoing silent transmission in specific hotspots, and highlights Moshi Rural as a persistently high-risk zone. Moreover, multivariable logistic regression showed that cattle in Moshi Rural had more than double the odds of seropositivity compared to those in Babati (p < 0.001), while no significant difference was observed between Hai and Babati (p = 0.657). Such spatial variation has been reported in previous studies and is often attributed to ecological and climatic factors, including vector density, altitude, and rainfall (Rugarabamu et al., 2021). Higher prevalence in Moshi Rural may reflect more favorable conditions for RVFV vectors, such as numerous water streams for irrigation on rice and sugar cane farms which cover an area of about 1100 hectares and presence of Nyumba ya Mungu dam which results to abundant population of Culex and Aedes spp (Rwegoshola *et al*., 2023).

No significant difference was observed between Rural (15.59%) and Peri-urban (14.01%) cattle systems (*p* = 0.424), this suggesting the similar exposure of risks factors across production systems, possibly due to shared grazing and watering points or unrestricted animal movement, which have been highlighted in other studies (Sumaye *et al*., 2013b; Sindato *et al*., 2015).

Sex as the predictor showed that female cattle had significantly higher odds of seropositivity than male, this has been also reported in other studies (Sindato *et al*., 2015; Salekwa *et al*., 2019) possibly due to longevity and herd retention hence more exposed to vectors.

Older cattle had significantly higher odds of exposure, other studies also identified older animals having higher odds compared young ones (Sindato *et al*., 2015; Umuhoza et al., 2017), supporting the hypothesis of cumulative exposure over time (Mohamed et al., 2010; Sumaye *et al*., 2013) This trend is consistent with the enzootic transmission of RVFV during inter-epizootic periods, where animals are silently exposed without overt clinical outbreaks.

Cattle from small herds had approximately four times lower odds of being seropositive compared to those from large herds, suggesting that larger herd sizes may facilitate greater exposure to vectors through generating more manure and moisture which attract mosquito breeding around animal enclosure, shared environments and also proximity of many cattle allow mosquito to feed across multiple animals. Access to water within crop fields increased the odds of seropositivity by 3.7 times, possibly due to increased mosquito breeding habitats in irrigated or wet environments, similar findings were reported by (Anyangu *et al*., 2010; Sumaye *et al*., 2013b).

Spatial analysis indicated that Seropositive cattle were not randomly distributed across the study area, but a clustering of RVFV exposure. Lower Moshi villages exhibited elevated levels of seropositivity compared to the other villages due to mosquitoes’ friendly habitats especially in irrigated Agro-pastoral areas. In Babati, villages located near Tarangire National Park and near Lake Manyara showed high number of seropositive cattle compare to other villages. These patterns point out the need for targeted surveillance and risk-based vaccination strategies, particularly in high-risk zones.

## 1.6 CONCLUSION AND RECOMMENDATION

This study demonstrates that there is ongoing inter-epizootic exposure of cattle to RVFV in northern Tanzania, forming distinct seropositive clusters rather than being randomly distributed. Higher seroprevalence was observed in Moshi compared to other study districts, while there were no differences of RVFV Exposure between rural and peri-urban settings. RVFV seropositivity clusters were observed in villages near key water sources, including Lake Manyara and Nyumba ya Mungu Dam, also villages near Tarangire National Park. Most significant determinants of seropositivity includes Age, Sex, Herd size, Water access along irrigated crop fields and geographical location Moshi showing higher seropositivity.

Future research should employ longitudinal designs with longer sampling periods to better capture temporal patterns of viral circulation. Sufficient resources should be allocated to include gold-standard confirmatory diagnostics, such as Virus Neutralization Test (VNT) or RT-PCR, alongside ELISA to improve accuracy and differentiate between past exposure and active infection. In addition, surveillance and control measures should prioritize identified hotspots and clustered areas rather than applying uniform strategies across all regions. Integrating spatial risk mapping into routine livestock health monitoring would further support timely and targeted interventions. Estimate the force of infection (FOI) using age-stratified seroprevalence data to quantify RVFV transmission intensity across different ecological settings. This will help distinguish areas with high ongoing transmission from those with lower exposure, guiding targeted surveillance and intervention strategies.

## CONFLICT OF INTEREST

There is no any conflict of interest on the issues presented in this article.

## ACKNOWLEDGEMENTS

The authors extend their sincere thanks to TVLA, SUA, KCRI, Agropastoral community, District Veterinary Officers, Livestock Field Officers, Livestock Officers, Ward and Village Executive officers, Local government authorities and farmers participated in Kilimanjaro and Manyara, for their cooperation and support to accomplishment of this study.

## APPENDICES

### Appendex 1 resarch approval from SUA

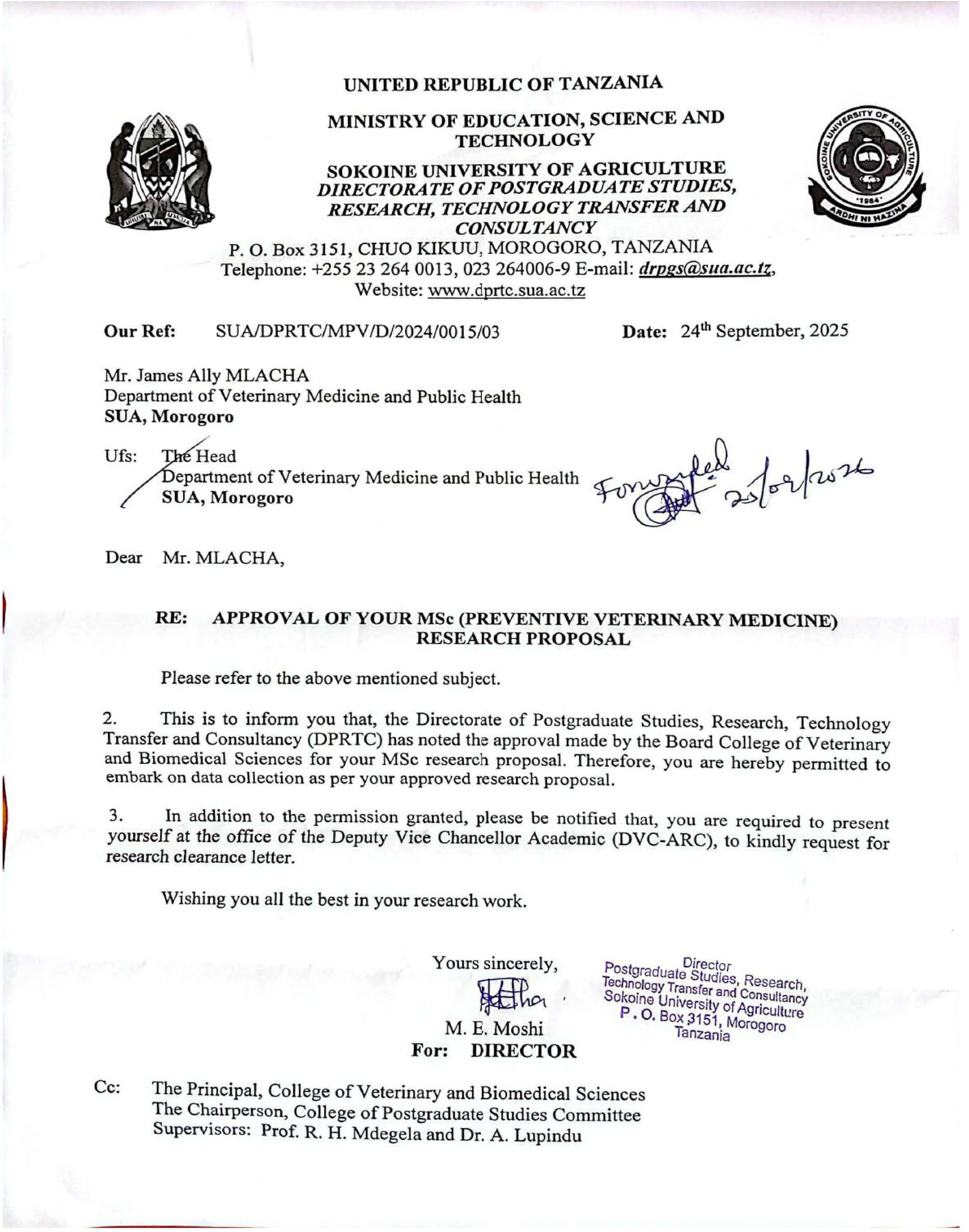

### Appendex 2 resarch approval from TALIRI

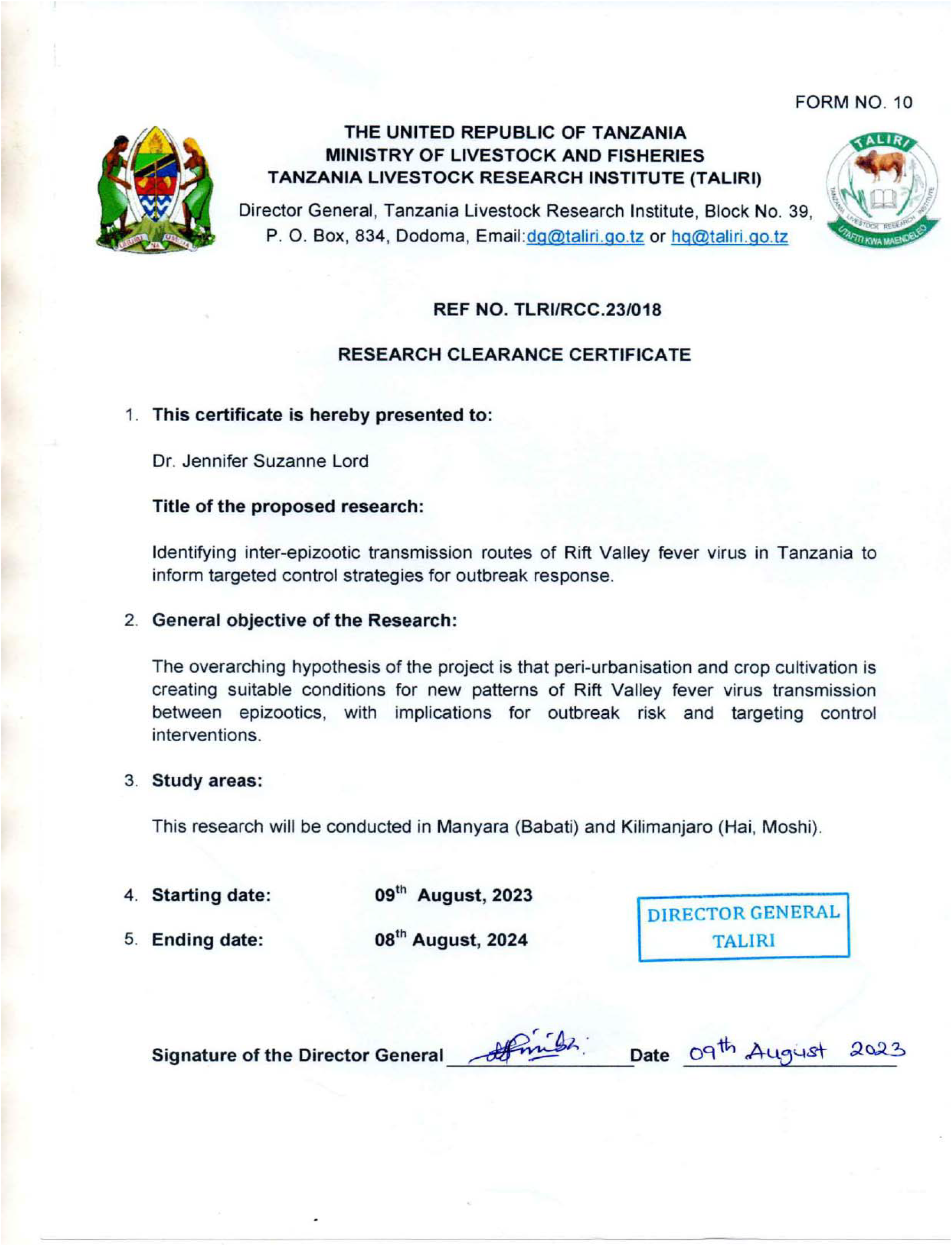

### Appendex 3 resarch approval from COSTECH

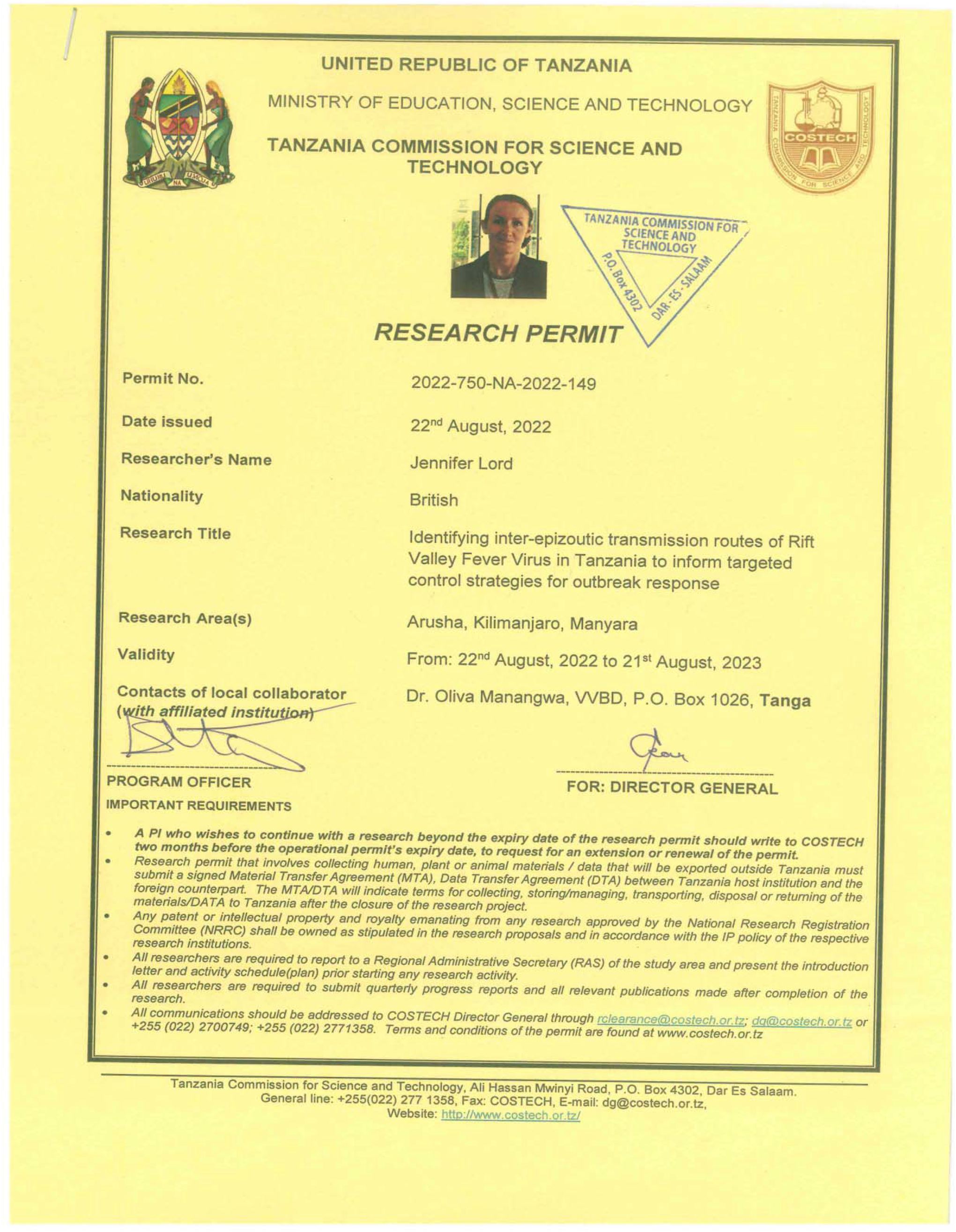

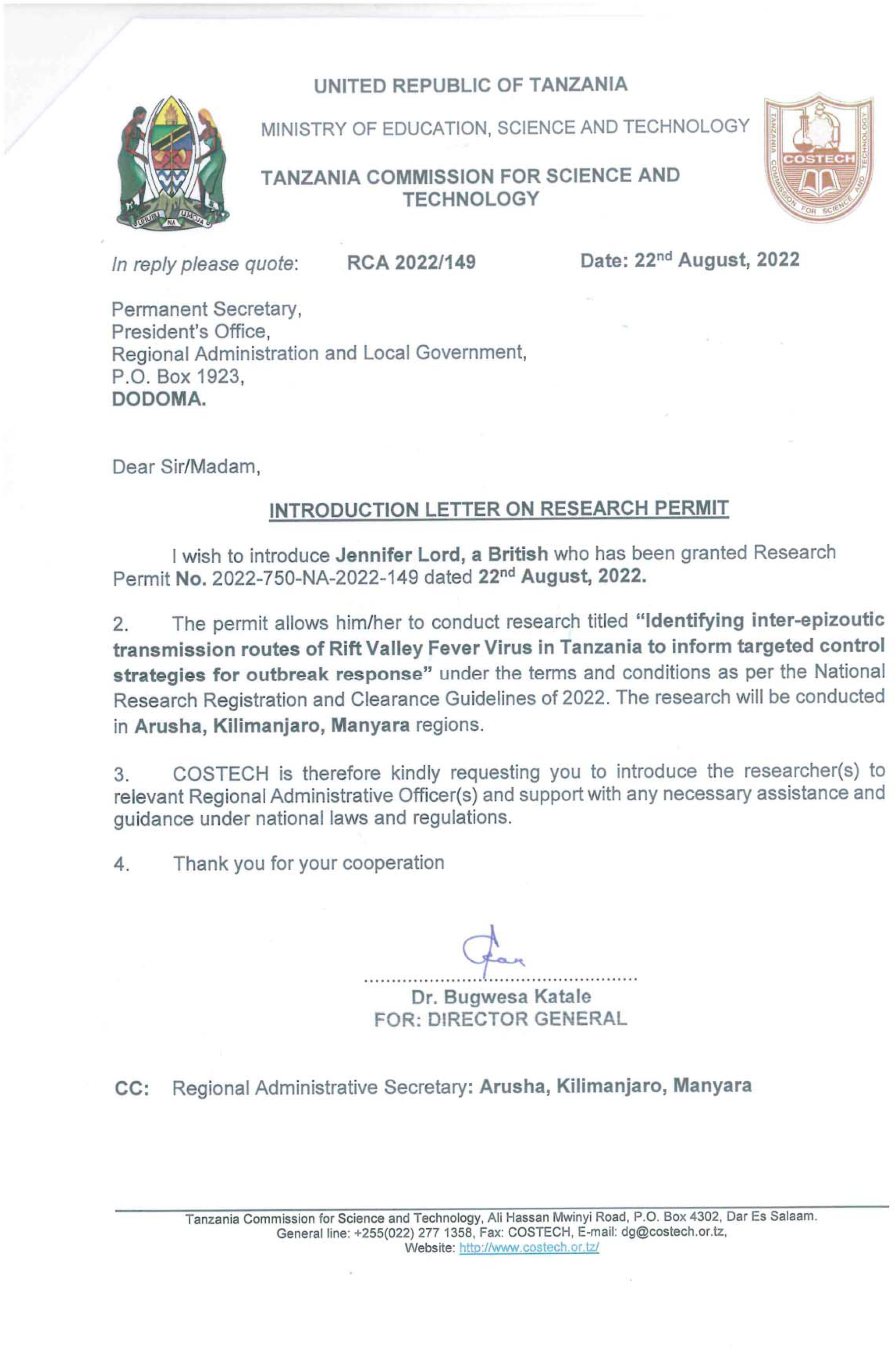

## Notes

### Competing Interest Statement

The authors have declared no competing interest.

### Funding

- Natural Environment Research Council: NE/W003333/1

